# Emerging arbovirosis (Dengue, Chikungunya, Zika) in the Southeastern Mexico: influence of social and environmental determinants on knowledge and practices. A mixed method study

**DOI:** 10.1101/581603

**Authors:** R Causa, MA Luque-Fernandez, H Ochoa Díaz-López, A Dor, F Rodríguez, R Solís, AL Pacheco Soriano

**Author notes:** Corresponding author. (HODL).

## Abstract

**Background:** The incidence and geographical distribution of arboviruses is constantly increasing. The epidemiological patterns of the proliferation of viruses and their vectors (*Aedes aegypti* and *Aedes albopictus*) are associated with socio-environmental determinants, and are closely related to human habits, especially at the household level. The aim of this work is to analyze the influence of socio-environmental determinants on the knowledge and practices related to arboviruses and their transmission, among the residents of three communities on the southern border of Mexico.

**Methodology:** Between June 2017 and August 2018, our investigation covered a set of 149 households from three communities of Tapachula (Chiapas) and Villahermosa (Tabasco). We first conducted household surveys about knowledge and practices on arbovirosis. Then, we carried out direct observations of risk factors for vector proliferation at the domestic level, before and after exposing a part of the population to a cycle of community engagement prevention activities. Through semi-structured interviews, we also focused on the detection of environmental risk situations for vector breeding at the community level.

**Key results:** We found that most dwellings had an adequate knowledge about the origin and transmission of arboviruses, but only a minority of them also implemented appropriate practices. Higher education levels were associated with better prevention scores. The observations made after the cycle of community activities in Chiapas revealed a decrease in the accumulation of unprotected water deposits. A higher percentage of domestic risk practices were detected in association with significant deficiencies in sanitation and water supply services. Furthermore, the perception of greater risk and difficulty in complying with preventive measures was detected among the population.

**Discussion:** Knowledge does not necessarily lead to adequate prevention practices. A better understanding of all these dimensions and their interactions is required. In addition to the educational level, intermediate social determinants (such as water supply and environmental sanitation) influence the persistence of behaviors that are risk factors for the proliferation of arbovirosis. The achievement of an effective and sustainable vector management is required to address these related aspects.

**Author summary:** Dengue, Chikungunya and Zika are arboviral diseases, transmitted by *Aedes* mosquitoes. As a result of a continuous increase in the geographical spread and burden of disease, arbovirosis have become a priority issue for global health.

The proliferation of viruses and their vectors are influenced by a complex interaction of environmental and social determinants. Thus, the design of effective and sustainable prevention and control measures requires an understanding of all these different aspects.

The aim of our work is to explore the effects of social and environmental factors on the knowledge and practices related to Dengue, Chikungunya and Zika and their transmission, among the residents of three communities on the southern border of Mexico, currently an endemic area. Our study draws on the application of a program combining the implementation of new technologies for vector management with a participatory and holistic multidisciplinary approach.

Between June 2017 and August 2018, we used different surveys and methodological approaches to explore knowledge and practices on arbovirosis, as well as to identify risk factors for vector proliferation. We found that intermediate social determinants (such as occasional water supply and infrequent waste collections) influence the persistence of behaviors that are risk factors for the proliferation of arbovirosis.

Public health interventions for arbovirosis prevention must be accompanied by intersectoral work that includes the improvement of these related aspects, according to the multifactorial etiology of arboviruses.

## Introduction

Dengue (DENV), Chikungunya (CHIKV) and Zika (ZIKV) are arthropod-borne viruses (arboviruses). In America, *Aedes aegypti*, is the main vector involved in its transmission. It’s a tropical mosquito of urban distribution, widely adapted to domestic environments, and with a worldwide distribution [1, 2].

In the context of the Region of the Americas, already endemic for DENV [3] and with high infestation levels of *Aedes* [2], the recent spread of CHIKV and ZIKV (with the first cases of autochthonous transmission at the end of 2013 and in the 2014, respectively) represent a threat and an important challenge for public health systems in the Region of the Americas [4-6].

The global burden of arboviral disease, and its rapid geographical spread, represents a growing public health problem and needs the formulation of effective strategies for the reduction of proliferation of viruses and their vector. Therefore, it is extremely important to understand the nature and interaction of the risk factors behind the emergence, reemergence and persistence of arboviral disease [1, 2, 7, 8].

The epidemiological patterns associated with the arboviral infection are the result of deep and complex interactions between biological, ecological, social, cultural and economic factors. Vulnerable contexts, with limited conditions of socio-economic development, weak health care systems and inadequate infrastructures are most disproportionately impacted by the infection, with consequences for the health and the economy that hinder the progress of the communities [7-12].

Human habits, especially at the domestic level, play a fundamental role in the spread of the disease, due to its close relationship with domestic and peri-domestic *Aedes* larval infestation indices: entomological indicator of the proliferation of the vector, and therefore, transmission risk of the disease [13, 14].

Reducing the vector population represent, currently, the only effective strategy to control arboviral transmission. Since the epidemiological patterns of the arboviral disease are associated with a set of social and environmental determinants, World Health Organization (WHO) strongly recommends the implementation of Integrated Vector Management as a required model for the control and prevention of arboviral disease [7-9, 15].

Our study draws on the application of a Multidisciplinary and Transversal (MT) project “Desarrollo de tecnología para el manejo integral de mosquitos vectores de dengue, chikungunya y Zika en Guatemala y México” (“Mosquito-MT project”), a prevention pilot program combining the implementation of the Sterile Insect Technique (SIT) with a participatory and holistic approach [16, 17]. Even if the SIT could represent an effective and sustainable alternative to traditional vector control programs, its introduction should be accompanied by an integrated approach including participation of local stakeholders. It has been remarked the importance of taking a community-based approach in vector borne disease interventions, especially when they include the implementation of new technologies for health [11, 18-21].

In recent years, the southern border of Mexico has been affected by repeated outbreaks and epidemics, with a high impact on the health and the economy of the area. It is currently a highly endemic area for the three diseases [22, 23].

The aim of this work is to analyze the influence of socio-environmental determinants on the knowledge and practices related to ZIKV, DENV and CHIKV and their transmission, among the residents of three communities on the southern border of Mexico, before and after carrying out a cycle of community activities with a part of the target population of the “Mosquito-MT project.”

## Methods

### Study setting and participants

The study was conducted between June 2017 and August 2018 at three rural villages on the southern border of Mexico, on the outskirts of two large urban centers: *Ejido* Hidalgo, *Ejido* Río Florido (near Tapachula, state of Chiapas) and *Ranchería* Guineo Segunda Sección (near Villahermosa, state of Tabasco).

According to the National Institute of Statistics and Geography, in 2010 the estimated population of Guineo Segunda Sección was 1032 inhabitants (298 dwellings), with 24.9% of the population over 15 years of age without complete elementary school. Besides, the estimated population of the Tapachula’s villages was 789 inhabitants (172 dwellings) in Rio Florido and 697 inhabitants (184 dwellings) in Hidalgo. The percentage of the population over 15 years of age without complete basic studies was 29.1% in Río Florido and 36.7% in Hidalgo. We gathered the obtained data of both villages because of their proximity (3 km apart), and the similarity of their overall condition [24].

Three different surveys were planned and the sample size for each was 149 households, with a total of 643 residents (82 households in Tapachula municipality, 67 households in Villahermosa municipality) and 6 key informants (2 participants from each community). The households were chosen through simple random sampling, while the key informants were chosen through purposive sampling. The key informant respondents were recruited primarily through the local community health center and the local community organization *Asamblea ejidal* [18]. A person from the *Comisariado ejidal* (representation of the *Asamblea ejidal)* of each community accompanied us during the process of the fieldwork, to achieve better understanding and acceptance with the rest of community.

In the Tapachula’s villages, two of the co-authors (AD and ALP) were conducting a community engagement plan, between November 2017 and July 2018. Through a collective and co-participative work, involving different social actors of the communities, different topics related to vectors and viruses were addressed, in order to improve capacities and informed decision-making about arbovirosis management and prevention. In addition to the active participation in the *Asamblea ejidal* meetings, a cycle of activities (talks, participatory workshops, community theater) for specific groups (women, children, adolescents) was designed.

### Study design

We conducted a mixed methods study, structured in two complementary phases. Phase I: in June 2017, prior to the start of the community engagement interventions, we visited 149 randomly selected households. We applied one Household Survey about knowledge and practices on arbovirosis and one Risk Observational Assessment about risk practices for vector proliferation at the domestic level, per dwelling unit. Phase II: during July and August 2018, we re-applied the Risk Observational Assessment to the same households of previous phase, after exposing a part of the population to a cycle of community engagement prevention activities. We also applied a Community Background Survey, focused on the detection of environmental risk situations predisposing to vector proliferation at the community level.

### Surveys

#### Household Survey

In 2017, through a structured, standardized and precoded questionnaire (S1.A File), we first obtained information about the sociodemographic characteristics of the household residents. Then we focused on basic knowledge on arbovirosis and its prevention. We first asked questions related to the origin and transmission of arbovirosis i.e. “how do you think is Dengue/Zika/Chikungunya transmitted?” (categorized as knowledge in the S1.A File).

Finally, we asked about the use of preventive measures i.e. “how do you avoid from mosquito bites?” and “how do you keep mosquitoes from breeding in your house?” (categorized as referred practices in the S1.A File). The interviews were conducted using the head of family as the informant.

#### Risk Observational Assessment

In 2017 and 2018, through the application of a structured, standardized and pre-coded household observational checklist (S1.A File), we collected information about risk factors for vector proliferation at the domestic level i.e., unprotected water deposits (vases, flowerpots, cans, tires, drinking troughs, buckets, tank, water tank, cistern etc.), accumulation of solid organic and inorganic waste and others environmental elements (debris, trash, plants, animals). The observations were made in the exterior space of the household, or *patio* (courtyard), where the kitchen, dinning room and rustic bathroom were generally located. Interior space was generally compounded by 1 to 2 bedrooms and a common space (for the television set). In the villages, the *patio* (courtyard) was then a fundamental extension of the house [30]. (These observations are categorized as observed practices in S1).

#### Community Background Survey

In 2018, through precoded and semi-structured questions (S1.B File) to household residents and key informants, respectively, we assessed the presence of risk factors for *Aedes* breeding in the community. The questions focused on the sanitation and hygiene services i.e. the existing facilities for waste collection, the source and frequency of water supply.

The semi-structured questions were used to triangulate the information recorded in the domiciliary visits, undertaking interviews with six key informants from each community (nursing staff of the local health center, *Presidente del Comisariado Ejidal, ejidal* committee of health). During the interview, we also collected qualitative data to explore risk perceptions and community’s expectations about arboviral diseases and their prevention.

#### Data management and analysis

The fieldwork of data collection was realized by trained staff and internal students from ECOSUR, Andalusian School of Public Health and Touro University of California. Information was collected through paper questionnaires. Quantitative data was entered (twice) into a database, using spreadsheets files. All data files were checked and cleaned separately by field supervisors. The data files of all study phases were first analyzed separately. Subsequently, they were merged and analyzed together.

Qualitative data from the community background survey were written as field notes, initially by hand, immediately during or directly after the interview process. The information was then organized, through a data collection template, according to the main themes explored (sanitation and water services; perceptions about arboviral disease risk factors and its prevention). Data was described and analyzed in order to identify common or recurrent patterns among the explored themes.

#### Knowledge, referred and observed practice score system development

In order to evaluate each section of the proposed surveys, we created indices (S1.A File) to score the knowledge and practices in question, as was already designed in several previous studies [26-30]. Questions related to knowledge only allowed a correct answer. Each correct answer was rated 1 point. Questions related to practices, allowed several correct answers, so that each interviewee had up to five possible options. Each correct answer was rated with 0.5 points. In any case, the wrong answers, or lack of answer, were rated with 0 points. Direct observations, made in the interior and exterior spaces of dwelling unit, focused on two aspects: presence of unprotected water deposits and detection of organic or inorganic solid waste. 1 point was assigned when these elements were absent (observed practices adequate), 0 points if not. A knowledge or practice was considered appropriate when the “overall score” obtained was equal or greater than 60% of the maximum expected score.

#### Statistical analysis

With the data collected, we describe the results with mean and proportions for continuous and categorical variables, respectively. Then, we performed univariate and bivariate categorical statistical analysis. Finally, we fitted a logistic regression model to explore social and environmental risk factors associated with the reported arbovirosis knowledge and practices. We derived the odds ratios (OR) and their respective 95% confidence intervals (CI) and assessed goodness of fit using the Hosmer and Lemeshow statistical test. For all the statistical analysis we used R version 3.2.1 (R Foundation for Statistical Computing, Vienna, Austria) and the R Commander package version 2.4-4.

#### Ethics statement

The research project proposal was approved by the internal review board of the XXXIII Master in Public Health of the Andalusian School of Public Health and the Research Ethics Committee of ECOSUR, with reference number CEI-006-2018. According to article 23 of the General Health Law for Health Research of the Ministry of Health of Mexico [31], due to the nature of the investigation, only verbal informed consent was applied.

To conduct the first part of the fieldwork (Household surveys and Risk observation assessment in June 2017), the authors from ECOSUR have received funding by the “Fondo de investigación científica y desarrollo tecnológico de El Colegio de la Frontera Sur (FID 784), in the context of the “Mosquito-MT project”. The second part of the fieldwork was undertaken as part of a Public Health Master’s Dissertation [32].

## Results

### Sociodemographic characteristics

Table 1 shows the study sample sociodemographic characteristics by locality of residence. In the majority of households the average number of inhabitants was of three of more people (82,5%). Overall, the most common type of family was “young”, represented by a mean age for the household composition members of less than 35 years old (55%). There were no significant differences in education level, sex and age distribution between the localities of the two municipalities.

**Table 1.** Sociodemographic characteristics. (Households Survey and Community Background Survey)

|  | Tapachula (n=82) |  | Villahermosa (n=67) |  | Global (n=149) |  |  |
| --- | --- | --- | --- | --- | --- | --- | --- |
|  | N | % | N | % | N | % | *p value* |
| **Residents per household** |  |  |  |  |  |  |  |
| 1-2 | 17 | 20.7 | 9 | 13.4 | 26 | 17.4 | *0.503* |
| 3-4 | 33 | 40.2 | 29 | 43.3 | 62 | 41.6 |  |
| >4 | 32 | 39.0 | 29 | 43.3 | 61 | 40.9 |  |
| **Women per household** |  |  |  |  |  |  |  |
| 0-1 | 28 | 34.1 | 20 | 29.9 | 48 | 32.2 | *0.843* |
| 2-3 | 41 | 50.0 | 35 | 52.2 | 76 | 51.0 |  |
| >3 | 13 | 15.9 | 12 | 17.9 | 25 | 16.8 |  |
| **Average of women per houseld** |  |  |  |  |  |  |  |
| <50% | 58 | 70.7 | 48 | 71.6 | 106 | 71.1 | *0.902* |
| ≥ 50 % | 24 | 29.3 | 19 | 28.4 | 43 | 28.9 |  |
| **Family typology (average age)** |  |  |  |  |  |  |  |
| Young (<35 años) | 40 | 48.8 | 42 | 62.7 | 82 | 55 | *0.089* |
| Adult and old (>35 años) | 42 | 51.2 | 25 | 37.3 | 67 | 45 |  |
| **Average education (≥15 year old)** |  |  |  |  |  |  |  |
| Without schooling | 11 | 13.4 | 4 | 6.0 | 15 | 10.1 | *0.186* |
| Elementary incomplete | 17 | 20.7 | 12 | 17.9 | 29 | 19.5 |  |
| Elementary | 32 | 39.0 | 37 | 55.2 | 69 | 46.3 |  |
| Advanced (secondary and higher) | 22 | 26.8 | 14 | 20.9 | 36 | 24.2 |  |
| **Female average education (≥15 year old)** |  |  |  |  |  |  |  |
| Without formal education | 12 | 15.2 | 8 | 11.9 | 20 | 13.7 | *0.919* |
| Basic (primary) incomplete | 16 | 20.3 | 16 | 23.9 | 32 | 21.9 |  |
| Basic | 32 | 40.5 | 27 | 40.3 | 59 | 40.4 |  |
| Advanced (secondary and higher) | 19 | 24.1 | 16 | 23.9 | 35 | 24.0 |  |
| **Solid waste collection services** |  |  |  |  |  |  |  |
| Daily | 0 | 0 | 0 | 0 | 0 | 0 | *0.000* |
| Weekly | 0 | 0 | 65 | 97 | 65 | 43.3 |  |
| Monthly | 82 | 100 | 2 | 3 | 84 | 56.7 |  |
| **Water supply services** |  |  |  |  |  |  |  |
| Regular | 0 | 0 | 0 | 0 | 0 | 0 | *0.000* |
| Irregular | 0 | 0 | 54 | 80.6 | 54 | 36 |  |
| Absent | 82 | 100 | 13 | 19.4 | 95 | 64 |  |
| **Drainage system** |  |  |  |  |  |  |  |
| Present | 0 | 0 | 0 | 0 | 0 | 0 | *NA* |
| Partial | 0 | 0 | 0 | 0 | 0 | 0 |  |
| Absent | 82 | 100 | 67 | 100 | 149 | 100 |  |

### Community background contextualization

Significant difficulties in community sanitation were detected, since the frequency of solid waste collection by the municipal services was always monthly in Tapachula and mostly weekly (97%) in Villahermosa. We also found important problems in the water supply services, completely absent in Tapachula and mostly irregular (80.6%) in Villahermosa, as well as a complete absence of a public drainage system in both sites.

Both in Villahermosa and in Tapachula, the wells (private or communal) were indicated by all the informants as the main solution to solve the lack of piped water. In *Ejido* Hidalgo, the lack of a functioning community well, even forced those families without other resources to store rainwater.

Furthermore, a high level of discomfort and preoccupation was detected, especially with the issue of waste management, as reported by a member of the Rio Florido *Comisariado ejidal*: “Before, there were no such problems. Each family consumed what they had. Now everything comes in plastic, packaging, and then we do not know what to do with so many things. There is a payment landfill, but it is only for families that can afford it.” Three of the six informants defined the situation as “[institutional] abandonment”. In addition, four of them expressed the perception of an increased risk of infection for the community (“There are more and more mosquitoes” / “more and more people get sick”/ “this has already become a problem”), as well as the perception of greater difficulty in complying with preventive measures. As one member of health staff expressed: “The most important thing is to keep the *patios* clean. It is a task of the whole community, we all must do it to make it work. This is why it is very difficult.”

### Knowledge and referred vector control practices

Table 2 describes the scores about knowledge and practices, as determined by the Household survey. Almost all of the respondents had heard of the diseases (99.3%), and knowledge about arboviruses and their transmission was generally good (75.2%), especially in relation with DENV (81.2%). However, only in few cases (30.7%) adequate personal (how to protect themselves from mosquito bites) or home (how to avoid mosquitoes breeding in and around houses) prevention measures were reported.

**Table 2.**
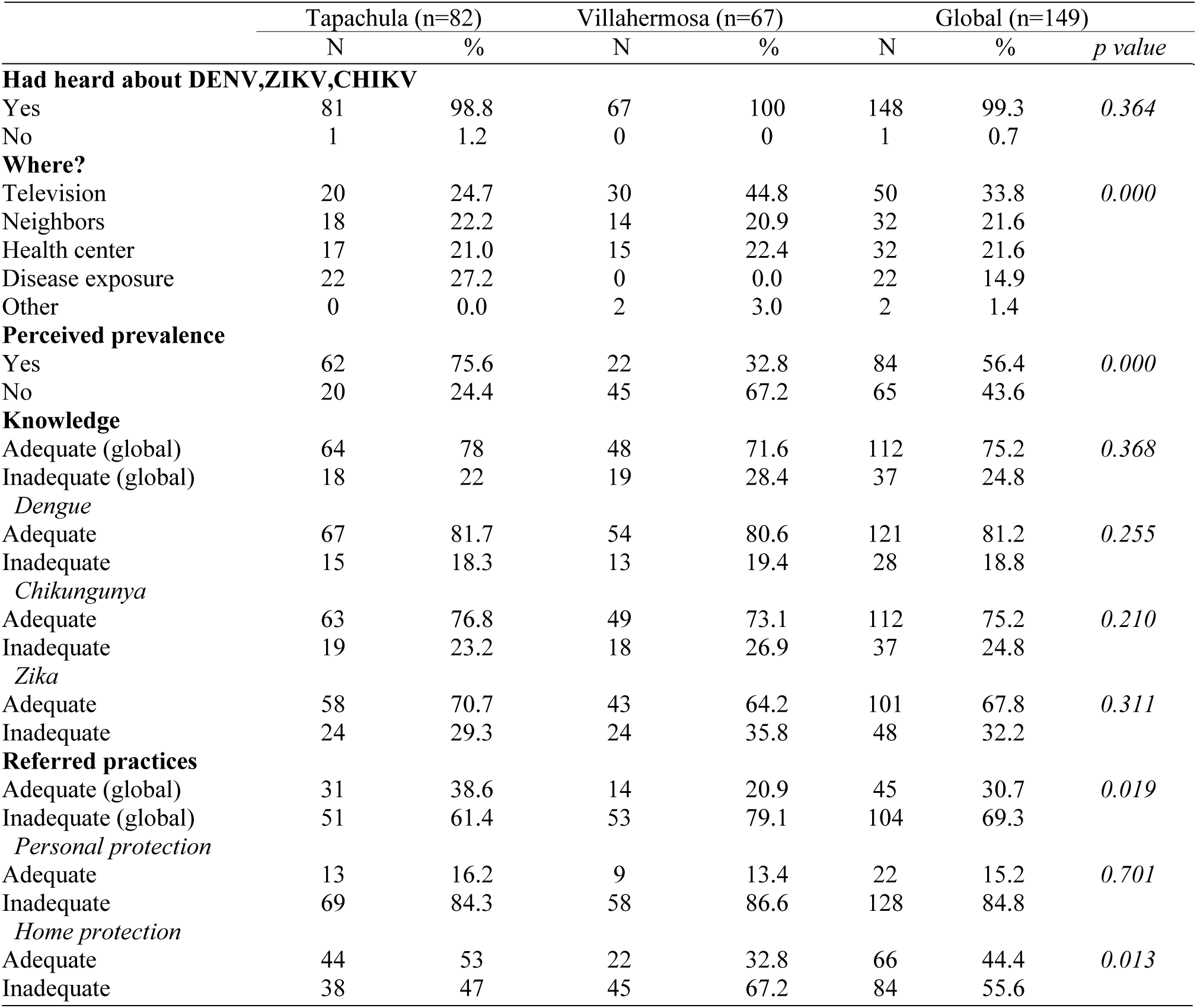
Knowledge and referred practices. (Household Survey)

### Observed practices

As Table 3 shows, both in 2017 and in 2018, a high proportion of households was found to have some unprotected water containers in outdoors areas (Tapachula: 73.7% and 42.1%; Villahermosa: 58.2% and 54.5% respectively), as well as waste accumulation (Tapachula: 60.5% and 65.8%; Villahermosa: 36.4% and 30.3% respectively). In 2017, Villahermosa’s village showed a higher percentage of adequate practices than in Tapachula’s villages (56.1% and 18.4%, p-value<0.001). However, in 2018, observed good practices showed an increase in Tapachula’s villages, from 18.4% to 27.6%. Besides, this change is completely associated with the decrease of unprotected water deposits, since the amount of organic and inorganic waste recorded in 2018 was even higher than in 2017. In Villahermosa’s village, the adequate observed practices slightly decreased from 56.1% to 42.4%, in relation with an increase of unprotected water deposits and waste in 2018. It is interesting to observe that the percentages of adequate practices in both sites are not significantly different in 2018 (p-value=0.064).

**Table 3.** Observed practices. (Risk Observational Assessment)

|  | 2017 |  |  |  |  | 2018 |  |  |  |  |
| --- | --- | --- | --- | --- | --- | --- | --- | --- | --- | --- |
|  | Tapachula,<br>(n=76) |  | Villahermosa,<br>(n=66) |  | *p-value* | Tapachula,<br>(n=76) |  | Villahermosa,<br>(n=66) |  | *p-value* |
|  | N | % | N | % |  | N | % | N | % |  |
| Adequate<sup>1</sup> | 14 | 18.4 | 37 | 56.1 | 0.000 | 21 | 27.6 | 28 | 42.4 | 0.064 |
| Inadequate<sup>2</sup> | 62 | 81.6 | 29 | 43.9 |  | 55 | 72.4 | 38 | 57.6 |  |
| *Unprotected water deposits* |  |  |  |  |  |  |  |  |  |  |
| Absent | 20 | 26.3 | 28 | 41.8 | 0.050 | 44 | 57.9 | 30 | 45.5 | 0.138 |
| Present | 56 | 73.7 | 39 | 58.2 |  | 32 | 42.1 | 36 | 54.5 |  |
| *Solid waste accumulation* |  |  |  |  |  |  |  |  |  |  |
| Absent | 30 | 39.5 | 42 | 63.6 | 0.375 | 26 | 34.2 | 46 | 69.7 | 0.000 |
| Present | 46 | 60.5 | 24 | 36.4 |  | 50 | 65.8 | 20 | 30.3 |  |

<sup>1</sup>Adequate: absence of unprotected water and/or waste.
<sup>2</sup>Inadequate: presence of water and/or waste accumulation.

### Logistic Regression Model

Table 4 shows the results of logistic models for the association between respondent sociodemographic characteristics and reported knowledge and practices. Higher education levels were associated with better knowledge and prevention scores. Household with uncompleted studies had 4-times (OR: 4.13; 95% CI: 1.2-13.9) higher probability of inadequate knowledge about arboviruses compared with household with primary or high studies, as well as higher probability of inadequate observed practices, both in 2017 (OR: 38.33; 95% IC: 7.87-186.53) and 2018 (OR: 15.04; 95% IC: 3.89-58.05).

**Table 4.**
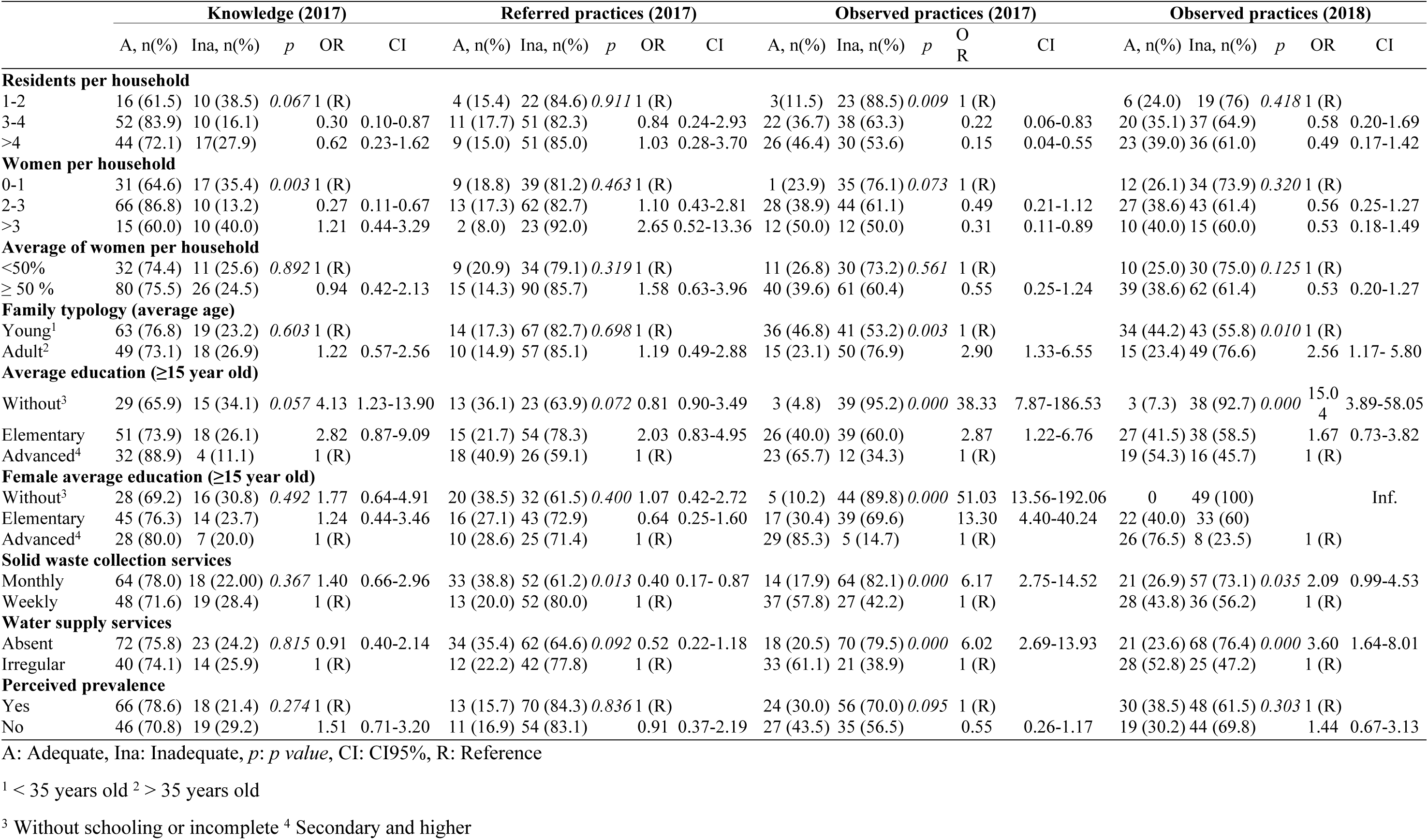
Respondent sociodemographic characteristics and reported knowledge and practices.

Likewise, a higher percentage of domestic risk practices were detected in association with significant deficiencies in sanitation and water supply services. When the frequency of solid waste collection was monthly, a higher risk of inappropriate observed practices was detected, compared to the weekly collection (2017: OR: 6.17, 95% CI: 2.75-14.52) (2018: OR: 2.09 95% CI: 0.99-4.53). Similar results were observed in relation to the absence of a water supply network, compared to its presence, even if irregular (2017: OR: 6.02, 95% CI: 2.69-13.93) (2018: OR: 3.60, 95% CI: 1.64-8.01).

## Discussion

### Principal findings and comparison with other studies

Reduction of *Aedes* vector population from the domestic environment is essential to control arboviruses transmission. It is basically achieved through the elimination of any larval habitat, since the domestic and peri-domestic *Aedes* infestation depends on their presence and quantity [13, 30]. Our findings suggest that social and environmental contexts influence the development of risk behaviors for vector breeding at the household level. In addition to the educational level, intermediate social determinants, such as water supply and environmental sanitation, are related with the persistence of risk factors for the proliferation of arboviruses.

Several studies evidenced the high impact of most of the socio-demographics determinants we described on domestic vector breeding risk [33-35]. In the context of Mexico, gaps in coverage of water and waste services was already identified as key factor for arbovirosis endemicity and the persistence of recurrent outbreaks [36, 37]. A study conducted in Cuba identified a significant association between irregularities in water supply and *Aedes* larval infestation indices [33]. Furthermore, outdoor water containers have been reported as the most productive vector breeding sites [38].

In our study, participants had high levels of knowledge regarding the transmission of DENV, CHIKV, ZIKV, but the overall score about self-reported and observed practices was generally low. Similar findings have been previously described in other localities with recent history of the disease [28, 29, 35, 39, 40]. These outcomes suggest that knowledge does not necessarily lead to adequate prevention practices, especially in those contexts where the complex interactions of environmental and social determinants increased human vulnerability to vector borne diseases [11, 12].

Community empowerment processes are a fundamental step to enable populations at risk (through behaviors and practices) to lead the eradication of vectors from their environment [15, 25]. There is previous evidence of the effectiveness of community-based approach strategies for the prevention of arboviral disease based on serological and entomological data [41-43]. The household risk observation assessment applied after the community engagement plan in Tapachula revealed a decrease in the accumulation of unprotected water deposits in the dwellings, as well as an increase in the accumulation of solid waste. Although it would be risky and inappropriate to propose a complete evaluation of the community engagement interventions, it seems that its potential impact, in particular on waste accumulation, could have been affected by contextual circumstances. These described circumstances were a high percentage of domestic risk practices detected in association with significant deficiencies in sanitation and water supply services. In Villahermosa’s village, where no community engagement plan has been operated, the sanitation services (water and waste collection) are more frequent than in Tapachula; and it is maybe why the residents of Villahermosa showed better practices than Tapachula’s residents in 2017.

In the semi-structured interviews, was recognized the importance of community involvement (especially at the household level) for the resolution of a problem perceived as growing, similar to the results of previous studies with a qualitative approach to the topic [44, 45].

Nevertheless, the difficulties expressed by the key members of the community during the interviews revealed the difficulty of translating preventive measures into everyday actions and behaviors.

### Limitations of the study

We are aware that our study instruments, chosen according to the available logistic and material resources, are not exempt from limitations. Surveys about knowledge and practices, due to their extension and quantitative approach, impeded a deep understanding of these dimensions and their interaction. Likewise, the categorization of survey data through a scoring system may not be the most appropriate method to approach such a complex issue, which is difficult to measure and quantify.

Furthermore, the reduced size and data sparsity regarding the surveys quantitative information possibly may have influenced a reduced statistical power for the statistical inferential analysis for some of the estimates associations.

### Conclusions and implications

We conclude that the endemicity of the arboviral infection in southern Mexico is the result of complex interactions between vector, environment and human behaviors. In order to develop targeted vector control programs to the southern Mexican area, a better understanding of the interactions between environment, vector and host human behaviors is needed. To achieve effective, sustainable, and ethical vector control strategies, a more comprehensive and contextualized approach is required. Community empowerment is more than just information: it is about facilitating and encouraging stakeholders to take control action, through strategies adapted to local conditions and needs. Furthermore, intermediary social determinants of water and sanitation need to be addressed to reduce community vulnerability to the arboviral disease [8-12].

## Acknowledgments

We would like to thank Alberto Fernandez, as well as the rest of the teaching team of the XXXIII Master in Public Health of the Andalusian School of Public Health, for providing academic and personal support to advance in this work.

We also acknowledge the staff of the ECOSUR Health Department of San Cristóbal de Las Casas and Villahermosa; especially, Rosario Garcia Miranda for the logistical support, Itandehui Castro Quezada for the statistical advice, and Cesar Irecta for his help in the field work in Villahermosa.

We are very grateful to the residents of Ejido Hidalgo, Rio Florido and Villahermosa, for all the support, contributions and time they have offered us.

## Supporting information

### S1 File – Surveys

**A. Household survey and Risk observational assessment.** Sections and simplified scoring system.

**B. Community background survey.** Adaptation of the pre-coded questions and semi-structured interview guide.

